# Allele age estimators designed for whole genome datasets show only a moderate reduction in performance when applied to whole exome datasets

**DOI:** 10.1101/2024.02.01.578465

**Authors:** Alyssa Pivirotto, Noah Peles, Jody Hey

## Abstract

Personalized genomics in the healthcare system is becoming increasingly accessible as the costs of sequencing decreases. With the increase in the number of genomes, larger numbers of rare variants are being discovered, leading to important initiatives in identifying the functional impacts in relation to disease phenotypes. One way to characterize these variants is to estimate the time the mutation entered the population. However, allele age estimators such as those implemented in the programs Relate, Genealogical Estimator of Variant Age (GEVA), and Runtc, were developed based on the assumption that datasets include the entire genome. We examined the performance of each of these estimators on simulated exome data under a neutral constant population size model, as well as under population expansion and background selection models. We found that each provides usable estimates of allele age from whole-exome datasets. Relate performs the best amongst all three estimators with Pearson coefficients of 0.83 and 0.73 (with respect to true simulated values, for neutral constant and expansion population model, respectively) with a 12 percent and 20 percent decrease in correlation between whole genome and whole exome estimations. Of the three estimators, Relate is best able to parallelize to yield quick results with little resources, however, Relate is currently only able to scale to thousands of samples making it unable to match the hundreds of thousands of samples being currently released. While more work is needed to expand the capabilities of current methods of estimating allele age, these methods show a modest decrease in performance in the estimation of the age of mutations.

**Article Summary:** Increasing availability of whole exome sequencing yields large numbers of rare variants that have direct impact on disease phenotypes. Many methods of identifying the functional impact of mutations exist including the estimation of the time a mutation entered a population. Popular methods of estimating this time assume whole genome data in the estimate of the allele age based on haplotypes. We simulated genome and exome data under a constant and expansion population demography model and found that there is a decrease in performance in all three methods on exome data of 15-30% depending on the method. Testing the robustness of the best performing method, Relate, further simulations introducing background selection and varying the sample size were also undertaken with similar results.

## Introduction

With the rapid advancement in sequencing technology and the availability of genomic data, such data has become integral to investigating the genetic cause of many major diseases. A major benefit of whole genome sequencing is the identification of mutations across the entire genome including non-coding regions, however it remains the costlier option in comparison with whole-exome sequencing (Warr et al. 2015; Schwarze et al. 2020; Almogy et al. 2022). Whole-exome sequencing (WES) targets the coding regions of the genome allowing for investigation into functional genomic mutations, which can more easily be associated with phenotypic changes and these mutations are found more commonly in databases of disease-causing variants (McLendon et al. 2008; Landrum et al. 2018; Karczewski et al. 2020; Piñero et al. 2020). This is more easily accomplished for monogenic diseases where just one gene contributes to a disease phenotype, however, improved methods have made understanding and identifying potential underlying mutations to polygenic diseases possible (Cano-Gamez and Trynka 2020). In particular, rare variants have been found to be associated with complex human diseases and phenotypes (Momozawa and Mizukami 2021) making variants at low frequency of particular interest to researchers.

As increasing numbers of genomes are sequenced, researchers are finding more rare variants, some of which have been shown to be causative of human disease (Yang et al. 2013). It is becoming more common for healthcare providers to perform exome sequencing to better understand the underlying cause of a patient’s condition (Goh and Choi 2012; Ma et al. 2019; Salfati et al. 2019). However, while these studies can identify potentially causative mutations through association studies, the history of these mutations are not as easily able to be examined except through large population studies (Grzymski et al. 2020). Currently, there are initiatives to expand current genomic resources with corresponding medical information (Carey et al. 2016; Grzymski et al. 2019; Backman et al. 2021). In the UK, researchers have released nearly 500,000 exome datasets corresponding to other participant information collected over the last fifteen years (Backman et al. 2021). Other similar biobanks operated primarily by healthcare systems are also utilizing genotype and exome data in their electronic health record collection (Carey et al. 2016; Grzymski et al. 2019) with the United States launching the “All of Us” initiative which was expanded to include 250,000 whole genome sequences (The “All of Us” Research Program 2019).

One way to better understand the history of a potential causative variant is to estimate when that mutation was introduced to a population. Combining the approximate age of an allele with allele frequency allows for a better understanding of the selective pressure the mutation has undergone (Slatkin 2000). Studies that attempt to understand the selective pressure on rare causative variants of disease rely on small whole-genome studies which focus on just a handful of individuals who carry a particular mutation of interest (Hateley et al. 2021). A previous study focused on candidate genes to utilize the absolute allele age to compare the ages of shared variants in the population to private variants only found in the probands (Wilfert et al. 2021), while a second study utilized relative allele age comparing the age of the candidate risk allele to other variants only shared between the two carriers (Hateley et al. 2021). In both cases, the studies utilized whole genome data limiting their analysis to just a small portion of all publicly available genotype data as many large population studies have worked to expand the database of WES sequences yielding very large datasets. One example of this is the gnomAD project which contains over 750,000 exomes contributing to about 90% of the total data represented (Chen et al. 2024).

In one previous study, exome data was used to estimate the age of alleles in a dataset of several thousand exomes leveraging a coalescent-based method with results indicating that most deleterious variants arose recently in human history (Fu et al. 2013). In more recent years, improved methods to estimate allele age have been published that yield more accurate estimates utilizing haplotype-based estimates of genealogies (Platt et al. 2019; Albers and McVean 2020) and ancestral recombination graphs (Rasmussen et al. 2014; Speidel et al. 2019; Wohns et al. 2022; Zhang et al. 2023). These more recent methods of allele age estimation have been shown to estimate the age of a mutation from WGS with relative accuracy, however little to no work has been done to establish accuracy of these methods when using WES data. The three methods included here, identified by the programs that implement them, are Relate, GEVA, and Runtc.

All three methods used here rely on mutation accumulation on shared haplotypes to accurately estimate the branch length of the branch containing the focal mutation. When estimating from WES data, the missing mutations in the non-coding regions will introduce increased variance in comparison with WGS due to there being less data utilized in the estimates. We may also expect there to be an increase in bias depending on the estimator’s underlying assumptions and methodology. For example, in the case of Runtc, the missing mutations would mean that the estimated shared haplotypes are much longer than the true haplotypes. This would then estimate a much shallower tree in comparison with its true deeper tree yielding an underestimation of allele age.

Relate provides estimates of allele age as part of an estimate of the ancestral recombination graph (ARG) and is one of several methods that do so (Rasmussen et al. 2014; Ignatieva et al. 2021; Wohns et al. 2022; Zhang et al. 2023). Relate was chosen for this analysis as it performed similarly in accuracy and scalability in comparison with other methods (Zhang et al. 2023). In the methodology for Relate (Speidel et al. 2019), the local genealogy underlying a focal site is reconstructed based on clustering from a distance matrix of mutation variation between haplotype pairs computed from a hidden Markov model (HMM). Then using a coalescent prior, Relate employs a Monte Carlo Markov Chain (MCMC) algorithm to estimate the time of the branch containing the focal mutation. Similarly, Genealogical Estimator of Variant Age (GEVA) (Albers and McVean 2020) also employs an HMM to identify shared and non-shared haplotype pairs, but then subsequently samples from those pairs before calculating the posterior estimate of the age of the mutation from a joint clock model leveraging recombination and mutation. Estimates of the time of first coalescence (t_c_) by the Runtc program (Platt et al. 2019) are based upon the maximum shared haplotype (msh) which can be bounded by either a recombination or mutation event. First coalescent time is used as a proxy for allele age and is estimated using the likelihood of the msh.

To assess performance, data sets were simulated under a constant population size model, a more complex demographic model considering the expansion of human populations out of Africa, and a model that included background selection. For each simulation, the true age of each mutation is compared to the estimates of the age generated from each program. Because rare variants are so integral to understanding human disease (Wang et al. 2021), we also examined specifically of how estimators performed on rare variants.

## Methods

### Simulated Datasets

Data sets were simulated with neutral mutations under both a simple, constant population model (Model id: PiecewiseConstant) and a complex demography model (Model id: OutOfAfrica_3G09) (Gutenkunst et al. 2009) using msprime version 1.2.0 (Kelleher et al. 2016) and stdpopsim version 0.2.0 (Adrion et al. 2020). Simulations were run to generate sequences that are a facsimile of human chromosome 22. They were 51megabases in length and were generated using the GRCh38 HapMap genetic map for chromosome 22 (The International HapMap Consortium et al. 2007). For the simple model, a cohort of 7242 genomes was sampled, and from the complex model, 7242 CEU genomes were sampled. Full parameters for each simulation model can be found in the supplement (Tables S1 and S2). From each simulation, segregating sites were output in VCF format using tskit version 0.5.0 (Kelleher et al. 2018) without any mispolarization or sequencing errors introduced. The output from the simulation will serve as the whole genome dataset for chromosome 22 for downstream analyses. Sites with multiple mutations were excluded from the analysis. True mutation ages are extracted from the simulated ancestral recombination graphs (ARGs) using the tree file. The entire simulation pipeline is described in Supplemental Figure 1.

### Conversion to Exome Data

For each model, the simulated sequences were filtered for the coding regions of chromosome 22 using a bed file of exonic regions generated from a bed file containing all autosome coding regions from the UK Biobank (available at: https://biobank.ndph.ox.ac.uk/showcase/refer.cgi?id=3803) (Backman et al. 2021). Using the BCFtools version 1.10.2-27-g9d66868 (Danecek et al. 2021) filter function, the VCF outputs from stdpopsim were filtered just for the segregating variants that fell within the regions in the supplied bed file. These filtered VCF files serve as the WES data in the analyses. Each of these original simulation runs are annotated as WGS or WES, and then either SIMPLE for constant population size or COMPLEX for the population expansion model hereafter. In total, four VCF files were prepared, one for each of the possible conditions of simple and complex demography, and exome and whole genome (identified as WES_Simple, WES_Complex, WGS_Simple, and WGS_Complex).

### Allele Age Estimates

#### Genealogical Estimate of Variant Age (GEVA) Estimates (Albers and McVean 2020)

The chromosome column of each of the four VCF files first were converted to number format using BCFtools (Danecek et al. 2021) as required by GEVA. Each VCF file was then converted to a binary file using the GRCh38 HapMap genetic map (The International HapMap Consortium et al. 2007) supplied from stdpopsim (Adrion et al. 2020) annotated with the additional map position column in centimorgans (cM). To run the estimator program for all sites of a frequency of 2 copies or more, a list of all sites (k>= 2) was extracted and split into batches of 300 variants due to memory constraints as recommended in the documentation. For the simple model, the mutation rate was 1e-8 and the effective population size was 10000. For the complex model, the mutation rate was 2.35e-8 and the effective population size was 7300, based on the values used for the simulations. Estimates from the Joint Clock model that passed the supplied heuristic filter were used. This includes the mode of the posterior distribution of the concordant and discordant sampled haplotype pairs. Conversion and estimator programs are available at https://github.com/pkalbers/geva.

#### Time of Coalescence (t_c_) Estimates (Platt et al. 2019)

Time of coalescence, t_c_, maximizes the likelihood of recombination and mutation events over the maximum shared haplotype, msh, as determined by the shared haplotypes tract surrounding a focal allele in a sample of genomes. However, in the case of missing data due to only sequencing the coding regions, the msh may start or end due to mutation or recombinational events in the intronic or intergenic regions. In the case of exome sequence data, the dataset is missing mutations between the exons. Since this data is missing, the end of the shared haplotype would not be identified until the next closest exon. To account for this we implemented a modified estimator in which the shared haplotype terminates at a random position between the exon where the haplotype was found to end and the next closest exon to the focal mutation. With modified msh value measured to include this random distance, the maximum likelihood estimator of t_c_ is calculated as originally described (Platt et al. 2019). Also following Platt et al. (2019) prior to estimating coalescent times of singletons (variants with just one copy in a population) VCF files were first phased for singletons by placing the mutation on the longer of the two possible haplotypes for each singleton in the sample.

The Runtc program was run with groups based on their k (copy frequency number) value using the flag --k-range to speed up the estimation pipeline. For the simple and complex models, the mutation rates and effective population size values were based on the values used for simulations, and the same as used for GEVA. The same genetic map as used for the simulations and other estimates for GRCh38 was used with the --map flag. This was done on all variants across the frequency spectrum. Phasing and estimator programs available at https://github.com/jaredgk/runtc.

#### Relate Estimates (Speidel et al. 2019)

Using Relate version 1.1.9, each of the four VCF files were converted to haps/sample format. Each analysis was run using 12 threads and Relate’s supplied parallelization script. For the simple and complex models, the mutation rates and effective population size values were based on the values used for simulations, and the same as used for GEVA. Additionally, for the complex model, the full pipeline was run to estimate the allele age based on branch lengths re-estimated after estimating the historical population size. The same genetic map as used for the simulations and other estimates for GRCh38 was used with the --map flag. Relate is available at https://myersgroup.github.io/relate/index.html.

### Statistics

For every combination of estimator and simulation, four statistics were calculated: Root Mean Square Log Error (RMSLE) between estimated and true ages, and bias, Pearson’s R, and Spearman’s R between log transformed estimated mutational ages and log transformed true ages calculated from the simulation.

### Comparisons across Sample Size

For both the simple and complex models described above, a spectrum of sampled genomes was extracted to compare accuracy across sample sizes. Sample sizes of 100, 500, 1000, 2500, 5000, 7500, 10000, and 15000 genomes were used. The same parameters as described above for each simple and complex model were used. Following the same pipeline as above (Figure S1), the Tree file for each simulation was extracted and converted to a VCF which was then filtered for exon sites. Using just Relate, allele ages for the WES versions of the simulation samples were estimated.

### Simulating sites with background selection

Following the pipeline described in Figure S1, stdpopsim version 0.2.0 (Adrion et al. 2020) was used to simulate mutations under background selection using SLiM version 4.1 (Haller et al. 2019) and the Gamma distribution of fitness effects based on Kim et al (Kim et al. 2017). Relate was then used to estimate the ages of mutations from both the original entire chromosome output from the simulation (WGS) and the filtered exome version (WES) with the same parameters as described above.

## Results

To better understand the accuracy of current methods of estimating allele age on whole-exome data, the ages of neutral mutations were simulated under both a simple, constant population size model and a more complex model. The SNP data directly generated from these simulations serve as the whole-genome dataset (WGS). For the simulated whole-exome data (WES), the generated simulated data was filtered based on the exome regions sequenced from the UK Biobank (Backman et al. 2021). For each of the simulations (simple and complex, WES and WGS), three estimators of the age of mutations were generated using the programs Relate (Speidel et al. 2019), GEVA(Albers and McVean 2020), and Runtc (Platt et al. 2019).

Each method provided estimates for a different subset of variants depending on the filtering mechanism implemented. Relate generates estimates for sites that pass the filtering threshold with all sites identified as non-mapping or flipped being excluded. The t_c_ estimator (implemented in Runtc) can be applied to all mutations, including those found just once (i.e. singletons). GEVA generates estimates for mutations of a frequency of 2 copies or more, and in addition required that some sites near the chromosome termini be excluded.

Relate estimated allele age the fastest, completing the estimation of allele age for the WES dataset in less than one hour for both demography models, and for the WGS in less than 12 hours for the simple model and around 13.5 hours for the complex model (Table S3). Both GEVA and Runtc took much longer as neither can easily parallelize. In the case of both estimators, sites can be separated into batches to allow for each batch to be run separately, however if forced to run sequentially both estimators would take on the magnitude of days to complete even for the much smaller WES dataset. Both GEVA and Relate (when run in parallel with multiple threads) required large amounts of memory and required dedicated computational resources.

### Relate Outperforms with a Simple, Constant Population Size Model of Neutral Variation

For each estimator method and simulated dataset, the estimated age of each variant was compared to the variants’ corresponding true age. The true mutation age was extracted from the tree files from the simulation output. For each comparison, bias, Pearson’s correlation, and Spearman’s correlation was calculated between the log transformed estimated ages and the log transformed true ages.

For mutations simulated under a simple model, a comparison of the true age to the estimated age from WES dataset showed a Pearson’s correlation coefficient of 0.83 for the Relate method with both GEVA and t_c_ with correlations of 0.79 and 0.57 respectively (Figure 1A, 1B, 1C). All three estimators underestimated mutations that were exceptionally old, with t_c_ having the largest discrepancy in allele age. As would be expected, each estimator did perform better estimating the ages of the mutations from WGS dataset with correlations of 0.94 for Relate (WES: 0.83), 0.92 for GEVA (WES: 0.79), and 0.86 for t_c_ (WES: 0.56) (Figure 1D, 1E, 1F). t_c_ saw the biggest shift in correlation in performance between WGS estimates and WES with a 33% drop in the Pearson’s correlation value (Figure 1C, 1F) with GEVA and Relate showing similar shifts in the Pearson’s correlation with decrease of 14% for GEVA and 12% for Relate between the estimates based on the WGS and WES datasets.

**Figure 1.**
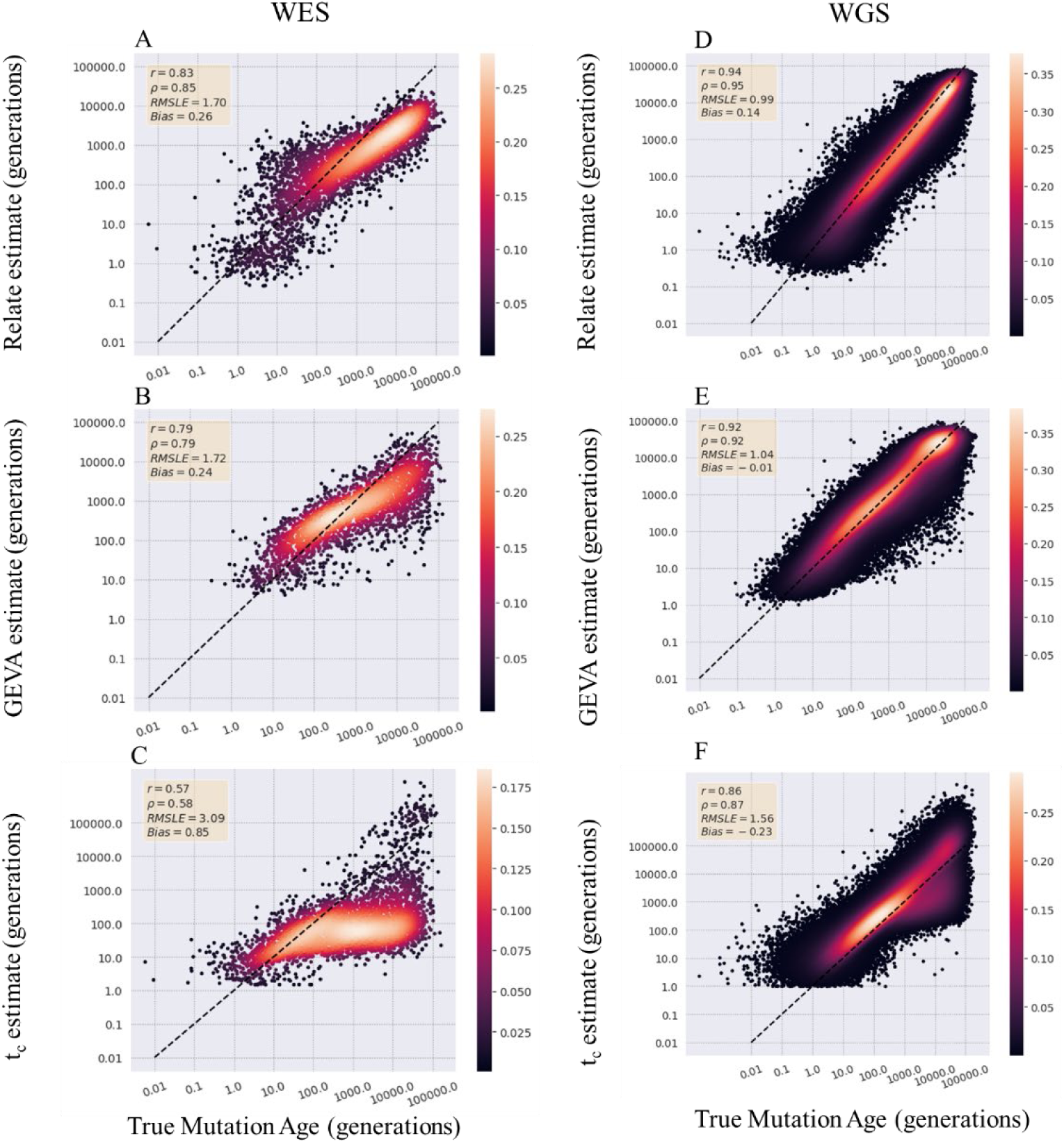
Estimator comparison on WES and WGS data simulated under a simple model. The estimated age of mutations was compared to true age values from variants simulated under a simple constant population size model. Values are reported estimates from WES datasets for Relate (A), GEVA (B), and time of coalescence (C) and from the WGS datasets for Relate (D), GEVA (E), and time of coalescence (F). Results are plotted on a log scale with the dotted black line representing perfect recapitulation of true age values. Points are colored by density calculated by a Gaussian density gradient. Pearson’s r, Spearman’s ρ, RMSLE, and Bias are reported for each comparison.

### All Three Estimators Show Relative Robustness to Demography and Selection

To investigate whether estimators of mutational age are robust to more complicated demography models, a more complex out-of-Africa model was simulated (Gutenkunst et al. 2009). Again, ages of all polymorphic mutations were estimated using the three estimators and compared with true values. A similar pattern of estimates from the simple model emerged with all three methods underestimating old mutations. Relate and GEVA show similar levels of correlation between the true and estimated ages with Pearson’s correlation coefficients of 0.73 for both with a lower correlation in the t_c_ estimated ages with a Pearson’s of 0.61 (Figure 2A, 2B, and 2C). Again, Relate and GEVA were better able to estimate the ages of the mutations from WGS data with correlations of 0.91 for Relate (WES: 0.73) and 0.86 for GEVA (WES: 0.73) (Figure 2D, 2E). Relate showed a 20% decrease in correlation between estimates from WGS and WES datasets while GEVA had a slightly smaller decrease in correlation at only 15%.

**Figure 2.**
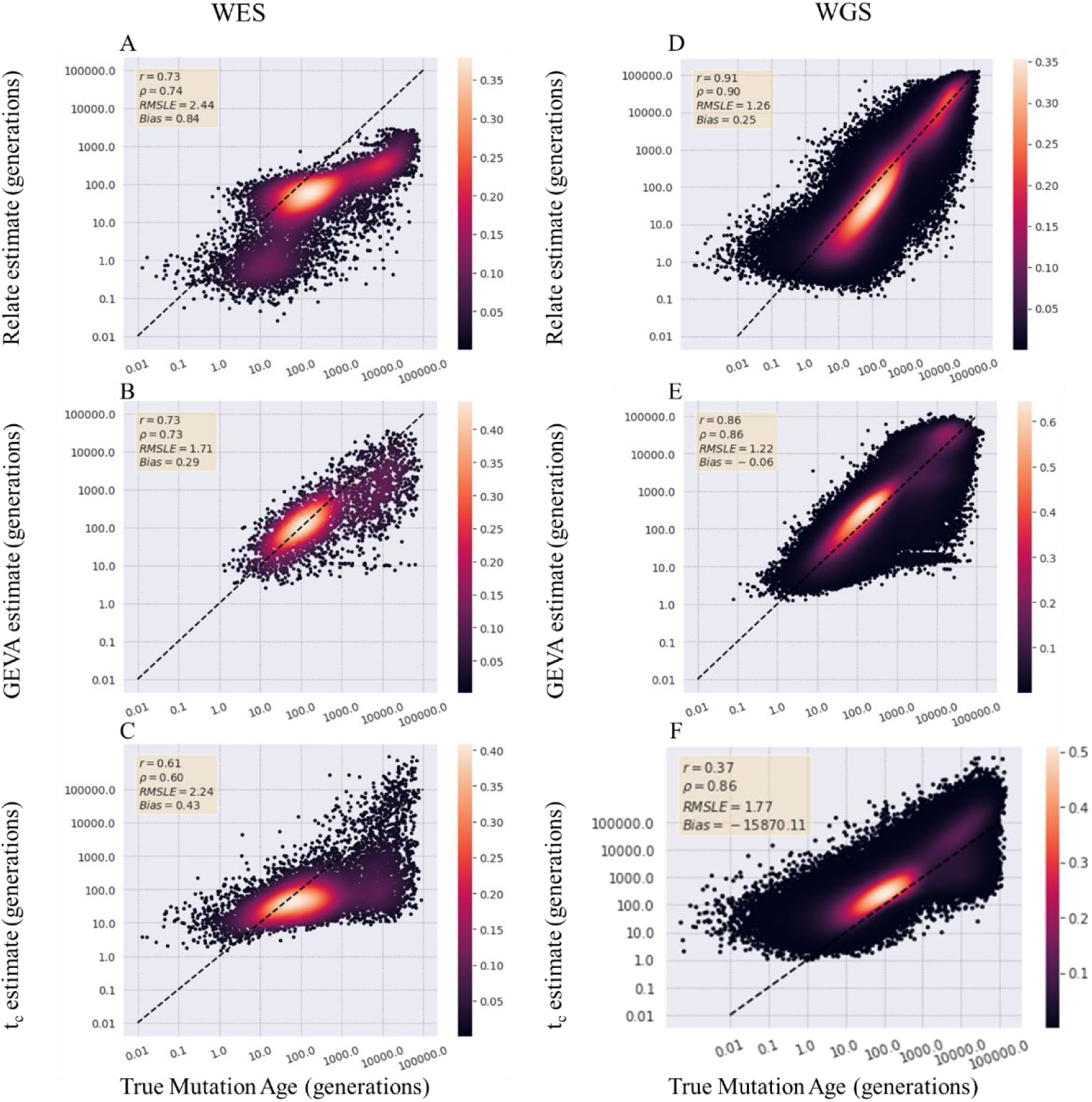
Estimator comparison on WES and WGS data simulated under a complex model. The estimated age of mutations was compared to true age values from variants simulated under a complex population expansion model. Values are reported estimates from WES datasets for Relate (A), GEVA (B), and time of coalescence (C) and from the WGS datasets for Relate (D), GEVA (E), and time of coalescence (F). Results are plotted on a log scale with the dotted black line representing perfect recapitulation of true age values. Points are colored by density calculated by a Gaussian density gradient. Pearson’s r, Spearman’s ρ, RMSLE, and Bias are reported for each comparison.

Further testing the robustness of allele age estimation on exome data, the estimator Relate was used to estimate the age of mutations in a sample size of 100, 500, 1000, 2500, 5000, 7500, 10000, and 15000 genomes. Relate was used as it was the most tractable for the larger sample size of 15000 genomes, however estimation of mutations in a sample of 100000 genomes was also attempted but required over 15 TB of storage and high memory requirements making it not easily tractable even with dedicated computing resources. It should be noted that an alternative ARG estimate, ARGNeedle, is designed to scale to hundreds of thousands of samples with similar accuracy to Relate and may be utilized with datasets of this magnitude (Zhang et al. 2023). With increasing sample size, the Pearson coefficient between estimated age of mutations and true age increased (Figure 3, Figure S2). This was previously reported for GEVA and Relate in which cases it was attributed to the increase in the surrounding mutations around a focal site to estimate the age of the mutation of interest.

**Figure 3.**
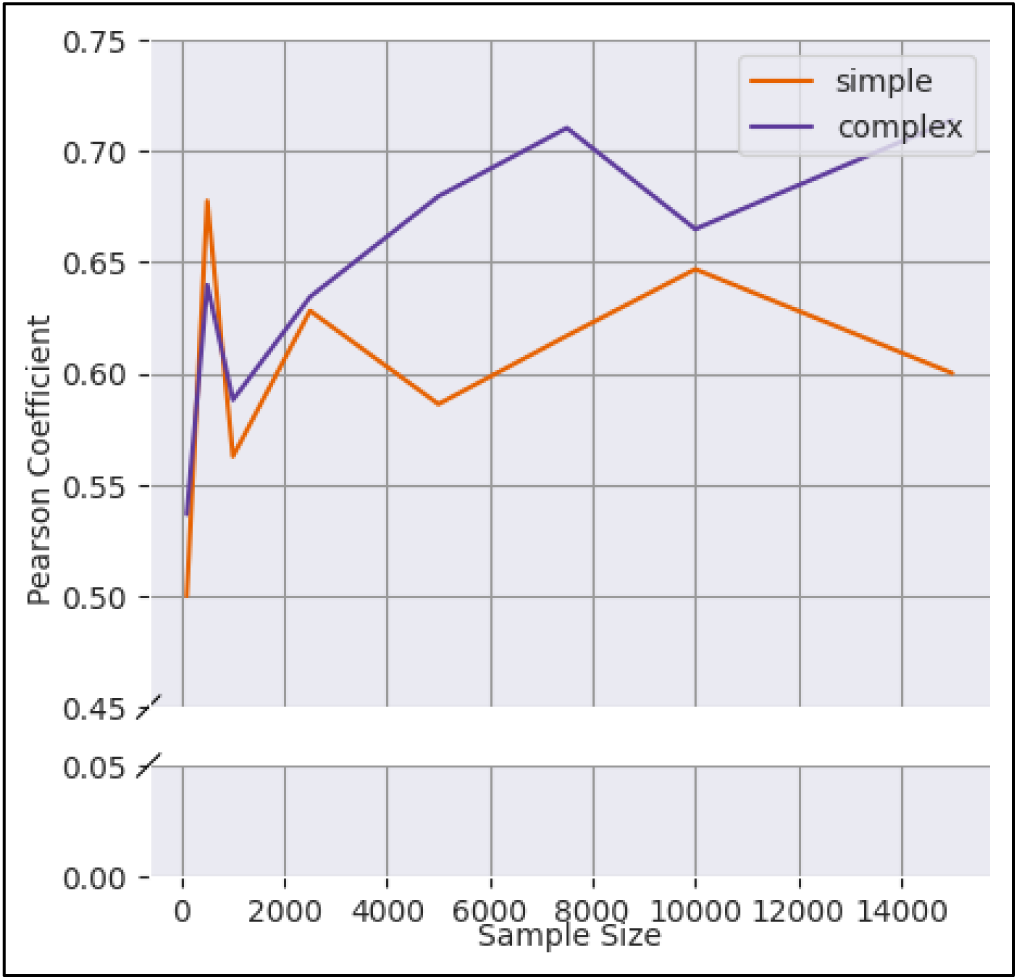
Correlation between true and estimated allele age increases with sample size. Allele ages are estimated from samples of 100, 500, 1000, 2500, 5000, 7500, 10000, and 15000 genomes from simulations of constant population size (purple) and population expansion model (orange). The true age of mutations is compared with the estimated age of mutations estimated with Relate for each of the three sampled set of mutations.

In nature, mutations are not just experiencing genetic drift, but often also natural selection, particularly negative selection because of a mutation being deleterious. To simulate this, keeping with the simple and complex model, each model is simulated using SLiM with background selection with a distribution of selection coefficients pulled from a gamma distribution. Relate was then used to estimate the ages of the mutations both using filtered exon dataset (WES) and the entire simulated chromosome (WGS). Relate performed slightly worse on data generated from a simulation with background selection, relative to its performance with no selection, for both the simple and complex models (Figure 4). Relate had an 18.6% decrease in Pearson’s correlation for mutations estimated from the WES dataset simulated under a simple model comparison with the WGS dataset. For mutations estimated under the complex model, there was an even higher decrease at 26.7% in Pearson’s correlation between the estimates. This is on average a 22% decrease in correlation for estimates under a non-neutral model, higher than the 16% average decrease in correlation between mutations estimated from datasets simulated with no selection.

**Figure 4.**
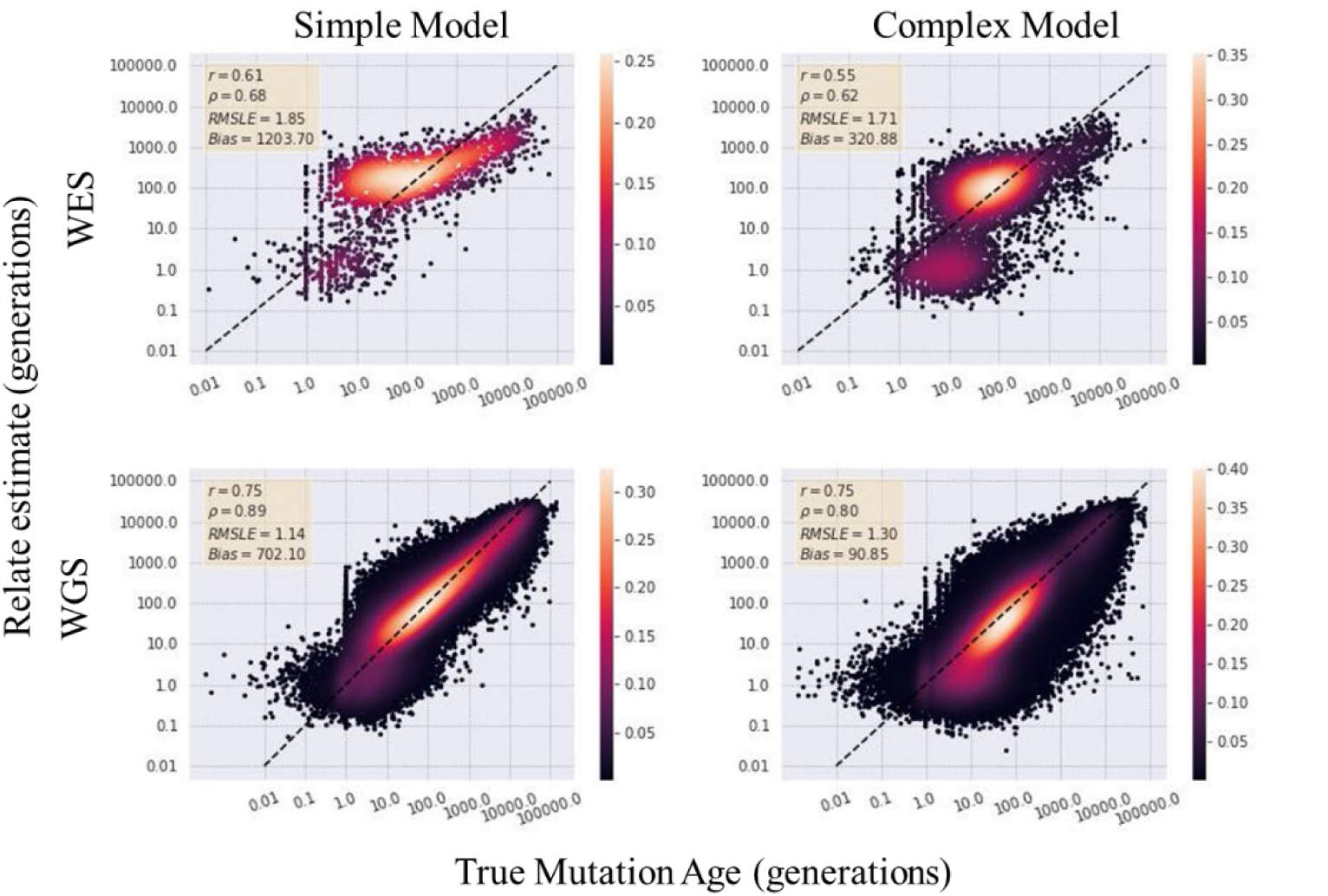
Relate allele age estimates compared to true values under background selection. The allele age of mutations from simulations of sampled genomes under a model which incorporates background selection are estimated using Relate. Upper panels have estimates from whole-exome datasets while lower panels contain estimates from the entire chromosome of mutations.

### Rare Variants Show No Decrease in Accuracy of Allele Age Estimates in Comparison with Common Variation

To address the accuracy of allele age estimators for rare alleles in WES datasets in comparison to more common variants, an analysis of the root mean square log error (RMSLE) averaged across frequency bins was undertaken. Variants were binned into 1% frequency bins and for each estimator method the RMSLE was calculated for that bin (Figure 5). GEVA and Relate had relatively similar RMSLE values for rare variants in comparison with common variants with larger amounts of variation between frequency bins for more common mutations (Figure 5A). However, time of coalescence showed much higher error rates for common variants (Figure 5A). Similar trends in error rates were found in the complex model (Figure S5). Focusing on rare variation, looking at the changes in error in mutations that just fall within the first 1 percent frequency bin, all three estimators show similar levels of error (Figure 5B). While GEVA is unable to estimate the age of singletons, both Relate and t_c_ were able to estimate the age of mutations appearing once in the population as well as other low frequency mutations (Figure 5B) with similar accuracy.

**Figure 5.**
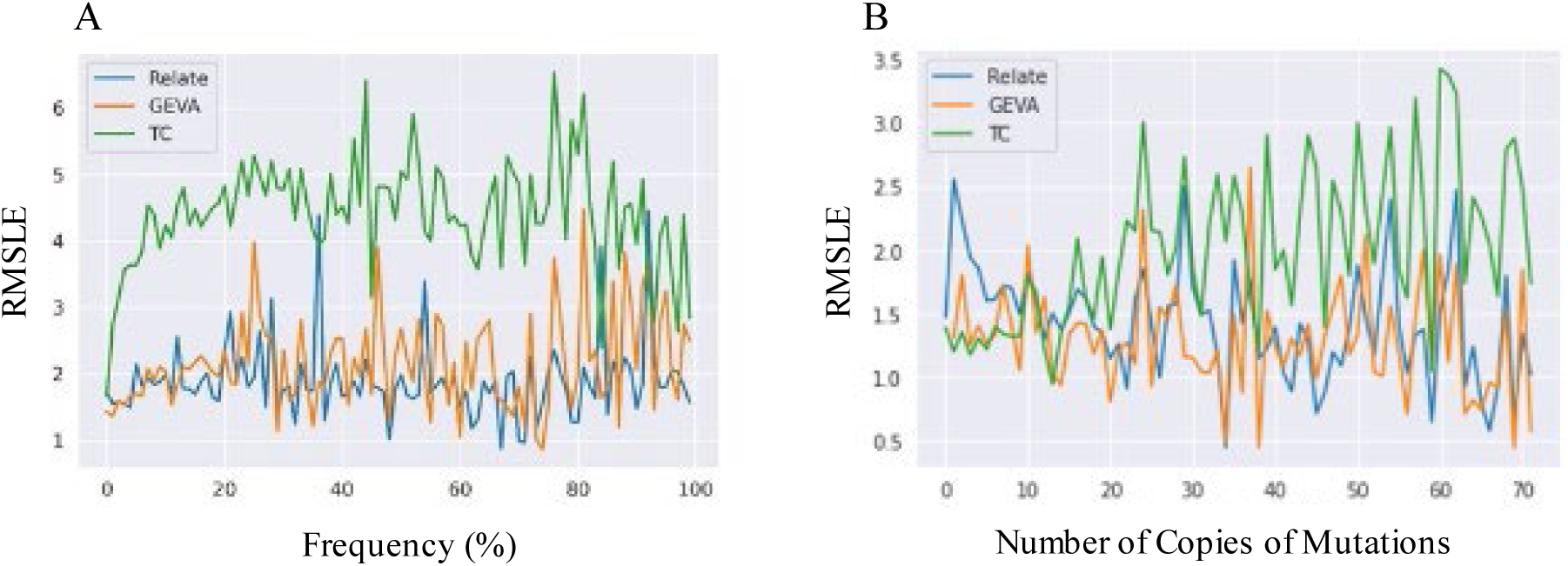
Error in estimates across spectrum of frequency values for the simple model. For each estimator, sites generated under the simple model are binned by 1% and an average root mean square log error (RMSLE) for that bin was normalized by the average true mutation age of that bin (A). Focusing on just k number of copies for 1% or less, the RMSLE is average for k values ranging from 0 to 72.

## Discussion

As sequencing becomes more cost efficient and thus more accessible (Schwarze et al. 2020; Almogy et al. 2022), personalized genomics will also become increasingly accessible. In personalized genomics, a large portion of resources is spent identifying and describing causative mutations of disease (Landrum et al. 2018; Karczewski et al. 2020), and in doing so, it becomes important to understand the functional impacts of the mutation. Currently, there are many methods to characterize a mutation (McLendon et al. 2008; Eilbeck et al. 2017; Landrum et al. 2018; Biddanda et al. 2020), with one approach being the estimation of the time of the mutation entered the population (Slatkin and Rannala 2000; Platt et al. 2019; Speidel et al. 2019; Albers and McVean 2020). As these methods are modeled and tested with whole genome sequence data little is known about their ability to estimate from whole exome data. As WES data remains the more cost-efficient option, it is more likely to be used when costs may be the prohibitive factor such as in personalized medicine (Goh and Choi 2012; Salfati et al. 2019). In one potential use case in characterizing a candidate variant of a disease phenotype, researchers estimated the ages of similar frequency variants to compare to the age of the candidate causative variant (Hateley et al. 2021). In studies such as this, allele age can be an additional indicator of the allele’s effect with harmful alleles having an expected age that is younger neutral alleles, conditional on frequency. However, it should be noted that these are noisy estimates and would only be useful in conjunction with other data. As age estimates can aid in understanding an allele’s effect especially in increasingly large datasets it’s important to identify the ability of current estimators to perform this task. Thus, here we conduct a comparison of several methods of allele age estimation to ascertain whether the methods can be successfully applied to whole exome sequence data.

Of the three estimators, as expected, all had the highest accuracy in identifying the ages of mutations from WGS data for both the simple and complex demography models with Relate performing the best across all estimators. For WES data, we find that of all three methods Relate again performs best with the highest correlation and lowest bias with a decrease of between 12 to 20 percent in correlation in comparison of the estimates on the WGS data. However, all three methods most underestimate the oldest of alleles. Time of coalescence underestimates allele age to the highest degree with the RMSLE tripling between 100 generations to 10000 generations. This indicates that the modified t_c_ estimator implemented here, that attempts to account for when a haplotype ends in non-coding regions, does not totally mitigate the issue of haplotype tracks ending in the intergenic or intronic regions. As a trend, each method slightly overestimated the youngest of alleles as the bias flips in signs from the youngest (less than 100 generations) to the oldest (greater than 100 generations). This is expected for older mutations, as coalescent events that happen further back in time allow for more mutations to accumulate along that branch. Thus, in estimating allele age from a dataset missing some substantial proportion of the mutations means that the estimators are observing fewer mutations on a branch and estimating it shorter than its true length. The overestimation of the youngest variants and underestimation of the oldest variants has been previously identified in comparison of whole genome sequence data estimates from both Relate and GEVA (Brandt et al. 2022; Ragsdale and Thornton 2023). In the comparison examining the robustness of the methods on simulations expanding beyond just a neutral model to also include deleterious mutations, Relate had a larger decrease in accuracy in the estimates between exome and genome data in comparison with the neutral model (simple: 19%, complex: 27%) (Figure 5).

All three estimators assume an infinite-site model where only a single mutation is able to arise at each site, however in nature this is not always the case (Karlin and McGregor 1967; Kimura 1969). For simulation data, this is not a concern as sites where a site had more than one mutation occur can be excluded or not allowed to occur. However, for empirical data, repeated mutations may occur especially in regions of the genome where the mutation rate is particularly high such as in CpG sites (Lek et al. 2016). If the infinite-site model is violated with repeated mutation at a given loci with multiple common ancestral haplotypes, then haplotypes with differing genealogical histories would then be pooled together artificially inflating the estimated ages. Relate deals with this issue by excluding sites where mutations cannot be mapped onto the local genealogy (Speidel et al. 2019) while t_c_ takes a composite of the estimated age of each copy of allele allowing for repeated mutations at a single locus not to skew the estimated ages too high (Platt et al. 2019). However, it is likely that the majority of human variant have arisen by way of single mutations, with violations of the infinite-site model by multiple mutations constituting a very small proportion of sites, given the few known examples (McVean et al. 2002; Blumenfeld and Patnaik 2004; Tishkoff et al. 2007).

While common variants have long been identified as potential pools of variants affecting human phenotypes (Burton et al. 2007; Claussnitzer et al. 2020), rare variants are often less well understood and harder to be studied and characterized. This is particularly important because these mutations are much less likely to be found in published large population data such as 1KGP (Auton et al. 2015) leading to very few opportunities to learn more about their history except through individualized studies or exceptionally large population studies. While rare variants have a lower error in comparison with common variants across all three estimators, it’s important to consider the range of ages for these low frequency variants. Common variants will have a wider range of variation in estimated values for alleles of the same frequency leading to higher error values. Thus, while the absolute age of a rare variant has a smaller error due to smaller variation, the relative position of the age of a rare variant within a given frequency bin may be more error prone to fluctuations in the estimators.

This analysis highlights both the abilities and limitations of utilizing allele age estimators on whole exome data to better understand the history of focal mutations. Expanding the methods of characterization for mutations of interest remains an important question as clinicians and researchers are still attempting to understand mechanisms of many disease phenotypes (Claussnitzer et al. 2020). These estimators demonstrate lower accuracy in absolute age estimates from whole exome data in comparison with whole genome estimates. However, all three estimators perform with similar accuracy on rare variants which introduces a potential opportunity to examine these previously hard to characterize variants. Relate which performs with the highest accuracy, has the potential to offer valuable insights into the history of potentially causative alleles to human disease through the comparison of relative ages among mutations of similar frequency. As the number of sequenced exomes increase with the expansion of personalized genomics, these methods can play a vital role in advancing our understanding of the genetic basis of disease.

## Supporting information

Full parameters for each simulation model can be found in the supplement

## Data Availability Statement

Simulations and code are available at https://github.com/ampivirotto/wes_alleleAge.

## Acknowledgments

We would like to thank Drs. Alexander Lucaci and Jordan Zehr for their thoughtful editorial insights.

