## Supplementary material for "Allele age estimators designed for whole genome datasets show only a moderate reduction in performance when applied to whole exome datasets": Full parameters for each simulation model can be found in the supplement

**Supplementary Information**

Table S1. Constant population simulation parameters

| **Parameter** | **Neutral Model** | **Selection Model** |
|---|---|---|
| Engine | msprime Version: 1.2.0 (Kelleher et al. 2016) | SLiM Version 4.1 (Haller et al. 2019) |
| Model | PiecewiseConstant<br>Piecewise constant size population model over multiple epochs. | |
| Seed | 14 | |
| Sample Size (genomes) | Varies depending on analysis: 50, 500, 3621, 5000 | 5000 |
| Contig Ploidy | 2 | |
| Sample Time (generation) | 0 | |
| Contig Length (bp) | 50818468.0 (Lander et al. 2001) | |
| Mean Recombination Rate (per bp per gen) | 2.1057233894035443e-08<br>(The International HapMap Consortium et al. 2007) | |
| Mean Mutation Rate (per bp per gen) | 1.29e-08 (Tian et al. 2019) | |
| Genetic Map | HapMapII_GRCh38<br>(The International HapMap Consortium et al. 2007) | |
| Generation Time (years) | 30 (Tremblay and Vézina 2000) | |
| Ancestral Population Size (individuals) | 10000 (Takahata 1993) | |
| Distribution of fitness effects (DFE) | | Gamma_K17<br>(Kim et al. 2017) |

The parameters for a simulation of a constant population model with either neutral mutations or negatively selected mutations. Citations listed for parameters. Documentation can be found at: https://popsim-consortium.github.io/stdpopsim-docs/stable/api.html#stdpopsim.PiecewiseConstantSize

Table S2. Complex population simulation parameters

<table>
<tr><th>Parameter</th><th>Neutral Model</th><th>Selection Model</th></tr>
<tr><td>Engine</td><td>msprime Version: 1.2.0<br>(Kelleher et al. 2016)</td><td>SLiM Version 4.1<br>(Haller et al. 2019)</td></tr>
<tr><td>Model</td><td colspan="2">OutOfAfrica_3G09<br>Three population out-of-Africa.<br>(Gutenkunst et al. 2009)</td></tr>
<tr><td>Seed</td><td colspan="2">14</td></tr>
<tr><td>Sample Size (genomes)</td><td>Varies depending on analysis: YRI:0; CEU: {50, 500, 3621, 5000}; CHB:0</td><td>5000</td></tr>
<tr><td>Contig Ploidy</td><td colspan="2">2</td></tr>
<tr><td>Sample Time (generation)</td><td colspan="2">0</td></tr>
<tr><td>Contig Length (bp)</td><td colspan="2">50818468.0 (Lander et al. 2001)</td></tr>
<tr><td>Mean Recombination Rate (per bp per gen)</td><td colspan="2">2.1057233894035443e-08<br>(The International HapMap Consortium et al. 2007)</td></tr>
<tr><td>Mean Mutation Rate (per bp per gen)</td><td colspan="2">2.35e-08</td></tr>
<tr><td>Genetic Map</td><td colspan="2">HapMapII_GRCh38<br>(The International HapMap Consortium et al. 2007)</td></tr>
<tr><td>Generation Time (years)</td><td colspan="2">25</td></tr>
<tr><td>Ancestral Population Size (individuals)</td><td colspan="2">7300</td></tr>
<tr><td>Distribution of fitness effects (DFE)</td><td></td><td>Gamma_K17<br>(Kim et al. 2017)</td></tr>
</table>

The parameters for simulation of an out-of-Africa population model based on Gutenkunst et al. 2009. Parameters indicated for either neutral mutations or negatively selected mutations.

Documentation for this simulation can be found at:

https://popsim-consortium.github.io/stdpopsim-docs/stable/catalog.html#sec_catalog_homsap_models_outofafrica_3g09

### Table S3. Timing of three estimators of allele age on exome and genome datasets.

| | Simple Model | | | | Complex Model | | | |
|---|---|---|---|---|---|---|---|---|
| | WES | | WGS | | WES | | WGS | |
| | Time (hrs) | Num SNPs | Time (hrs) | Num SNPs | Time (hrs) | Num SNPs | Time (hrs) | Num SNPs |
| Relate (12 threads) | 8.448 (0.704) | 3981 | 140.616 (11.718) | 171107 | 10.62 (0.885) | 9400 | 162.204 (13.517) | 407286 |
| GEVA | 18.62 | 2672 | 2110.3 | 120499 | 22.3 | 3254 | 1530.8 | 169149 |
| $t_c$ | 65.45 | 3965 | 3023.5 | 171876 | 82.52 | 9314 | 4474.8 | 406388 |

Each of the three allele age estimators (Relate, GEVA, tc) were run on neutral simulations of the simple constant population size model and the out-of-Africa expansion population size model for both the whole exome sequences (WES) and the whole genome sequences (WGS). The time each estimator took to return estimates was recorded in hours and the number of single nucleotide polymorphisms (SNPs) estimated for each dataset and method were also recorded. For Relate the actual time taken to run on 12 threads is recorded in the parentheses for each simulation model. Each estimator filtered some subset of the sites due to filtering mechanisms of the method as described in the methods.

Figure S1. Depiction of simulation pipeline.

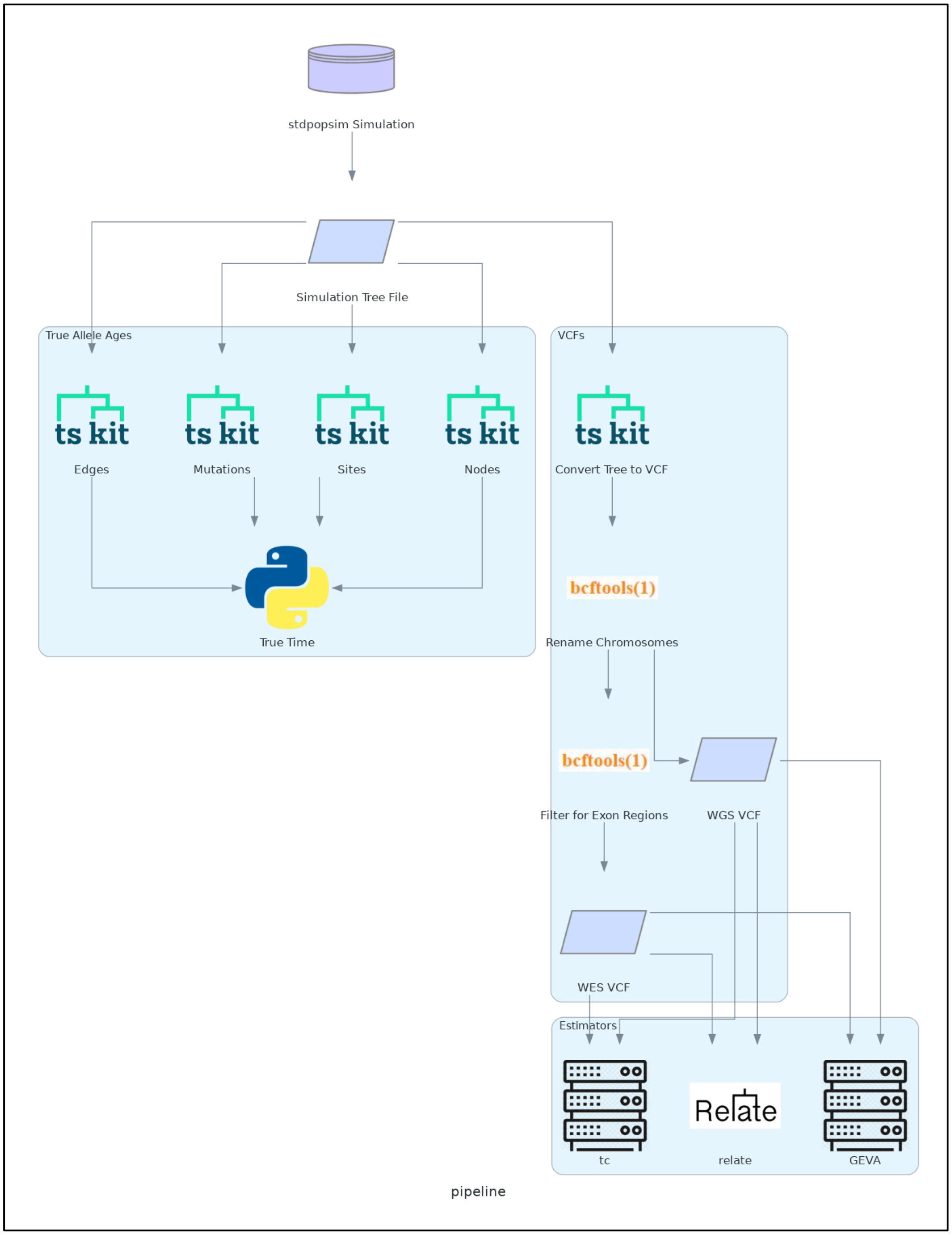

Pipeline for process of simulating dataset of mutations, extracting true age of mutations, converting simulations to VCF format and filtering for sites in the exon regions, and estimating age of sites using three estimators: GEVA, relate, tc.

24

### Figure S2. Comparison of Relate on samples of 100, 1000, and 10000 genomes.

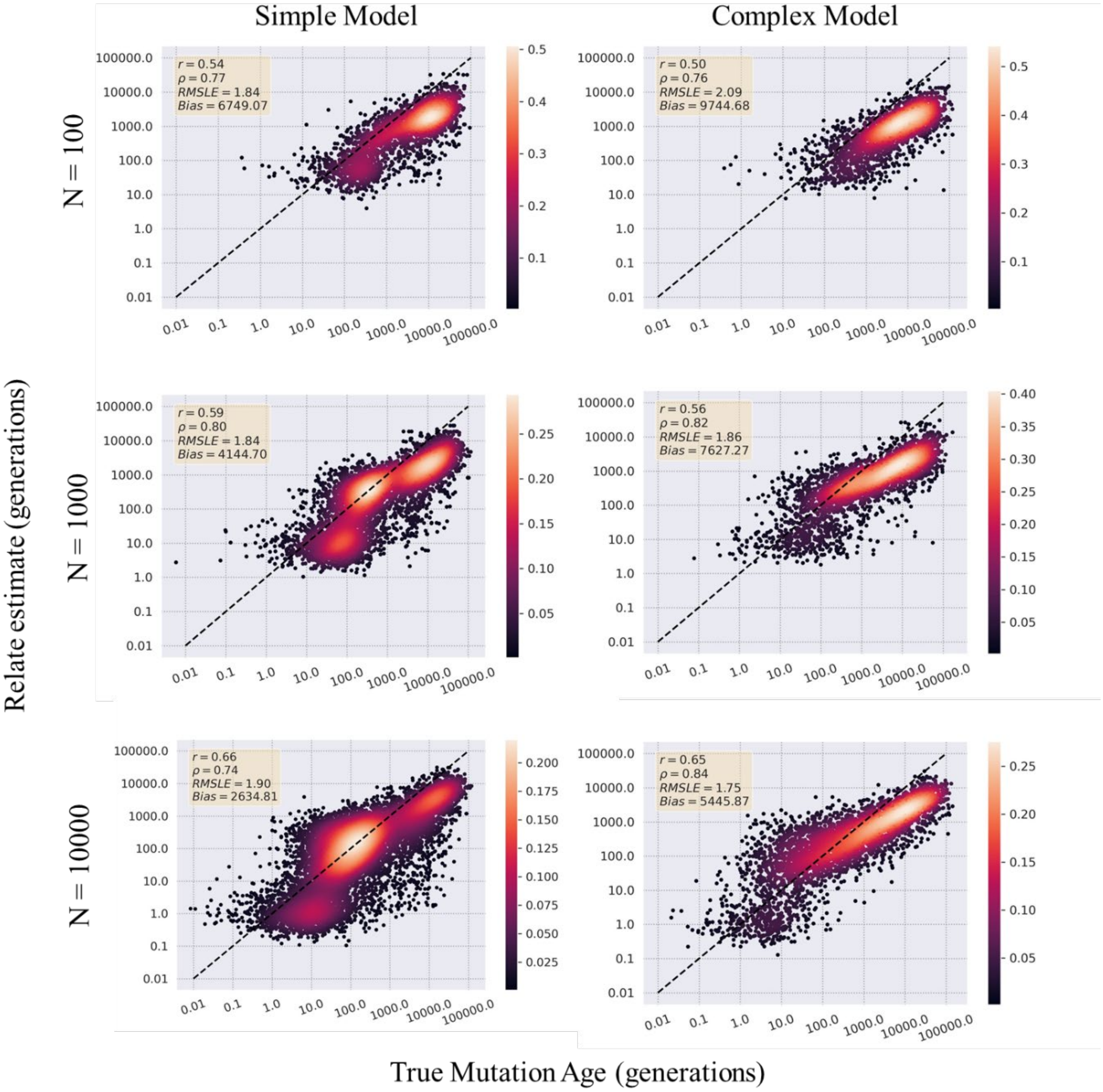

Neutral simulations under both a simple constant population size model and a complex out-of-Africa expansion population size model were undertaken with the sample sizes of 100 genomes (50 diploid individuals), 1000 genomes (500 diploid individuals), 10000 (5000 diploid individuals, and 100000 (50000 diploid individuals). Relate was run on the simulations of samples 100, 1000, and 10000 as the simulation with a sample of 100000 genomes was intractable due to large storage and time needs. Points are colored by the spatial density as calculated from a Gaussian kernel density. Pearson's r, Spearman's ρ, RMSLE, and Bias are reported for each comparison.
